# Characterization of antibiotic resistance development of *E. coli* in synthetic and real wastewater

**DOI:** 10.1101/2024.10.16.618744

**Authors:** Indorica Sutradhar, Neila Gross, Carly Ching, Yanina Nahum, Darash Desai, Devin A. Bowes, Muhammad H. Zaman

**Affiliations:** Department of Biomedical Engineering, Boston University, Boston, MA 02215, USA; Department of Materials Science and Engineering, Boston University, Boston, MA 02215, USA; Center for Forced Displacement, Boston University, Boston, MA 02215, USA; Department of Environmental Health Sciences, Arnold School of Public Health, University of South Carolina, Columbia, SC 29208, USA; Center on Emerging Infectious Diseases Research, Boston University, Boston, MA 02118, USA; Howard Hughes Medical Institute, Boston University, Boston, MA 02215, USA

## Abstract

Antimicrobial resistance (AMR) is a major threat to global health and resistant bacterial populations have been observed to develop and spread in and around wastewater. However, *in vitro* studies on AMR development are typically conducted in ideal media conditions which can differ in composition and nutrient density from wastewater. In this study, we compare the growth and AMR development of *E. coli* in standard LB broth to a synthetic wastewater recipe and autoclaved wastewater samples from the Massachusetts Water Resources Authority (MWRA). We found that synthetic wastewater and real wastewater samples both supported less bacterial growth compared to LB. Additionally, bacteria grown in synthetic wastewater and real wastewater samples had differing susceptibility to antibiotic pressure from Doxycycline, Ciprofloxacin, and Streptomycin. However, AMR development over time during continuous passaging under subinhibitory antibiotic pressure was similar in fold change across all media types. Thus, we find that while LB can act as a proxy for wastewater for AMR studies in *E. coli*, synthetic wastewater is a more accurate predictor of both *E.coli* growth and antibiotic resistance development. Moreover, we also show that antibiotic resistance can develop in real wastewater samples and components within wastewater likely have synergistic and antagonistic interactions with antibiotics.

**Importance:** Antimicrobial resistance (AMR) ranks among the leading global threats to public health and development. In 2019, bacterial AMR was estimated to have directly caused 1.27 million deaths worldwide and contributed to 4.95 million deaths overall (Murray, C. J., et al., (2022). Global burden of bacterial antimicrobial resistance in 2019: a systematic analysis. *The Lancet, 399*(10325), 629–655.). With estimations of AMR only getting worse, it is imperative that we understand the complex dimensionalities that drive the genesis of antimicrobial resistance to where it begins–the environment. The paper investigates bacterial growth and AMR in real wastewater samples and highlights the importance of using a media that closely mimics real wastewater in AMR studies, compared to standard lab media like LB broth. This is crucial for understanding how *E. coli* and other bacteria develop AMR in environments similar to actual wastewater, which can inform more effective strategies to combat AMR in natural and engineered settings.

## Introduction

Antibiotics and antimicrobials are essential medicines to treat infectious diseases. However, when bacteria gain antimicrobial resistance (AMR), these medicines no longer work, threatening humans, animals and the environment. Our understanding of the role of the environment in AMR is starting to increase. Wastewater is a key component of a larger and more complex shared environment in which AMR can spread and develop^1^.

Wastewater is a major reservoir of AMR due to the collection of antibiotic pollution from inappropriate drug disposal and effluent from pharmaceutical manufacturers, hospitals and agricultural/veterinary settings^2^. In some cases, environmental concentrations can reach, or even exceed, minimal inhibitory concentrations of certain^3^. This problem is particularly pertinent in LMICs where cases of antibiotic-resistant infections have been rising and 70% of sewage produced is estimated to enter the environment untreated^2^. Antibiotic pollution is a major driver of resistance, as selective pressure from antibiotics in the environment is known to promote chromosomal resistance mutations^3, 4^. These antibiotics, as well as disinfectants in sewage, may also be able to support and promote horizontal transfer of resistance genes among bacteria^5^ and such mobile resistance genes have been measured at high levels in wastewater and sewage^6^. Though AMR and its drivers have been studied in sewage and wastewater settings to some extent^7,8^, one of the largest gaps in understanding the emergence of AMR within a sewage environment is the limited understanding of the effects and interactions of the many biological and environmental mechanisms at work due to the complexity of wastewater as a matrix^9^.

Aside from antibiotics, wastewater is a complex matrix that contains a mixture of a variety of substances as a result of human, animal, and plant activity within the built and natural environments. Composition changes depending on the immediate local conditions that determine the sources of the constituents. However, generally all wastewaters contain water, organic matter, organic chemicals, nutrients, metals and other inorganic materials, and microorganisms (fungi, bacteria, protozoa, viruses, etc.)^10^. Wastewater treatment plants are widely recognized as potential hotspots of conferring antibiotic resistance due to this inherent complex mixture. Additionally, other factors such as population growth, increased use and distribution of antibiotics, and products from human activity such as metal-based pesticides, fertilizer runoff from agricultural practices, and persistence of toxic chemicals from personal care products all contribute to promoting favorable conditions for the growth and transfer of resistance genes within the generalized wastewater treatment infrastructure^11^. For instance, the use of pesticides, while pertinent for the production of food, has demonstrated to influence antibiotic resistance as certain types have shown to present with similar inhibitory mechanisms to some antibiotics, such as chlorhexidine and fenticlor^12^. Increased and continuous contamination of heavy metals from various industrial activities that runoff into the surrounding environment and ultimately result in municipal wastewater streams has encouraged microorganisms to evolve and develop co-resistance with antibiotic resistance. This has become of particular concern from a global health perspective in regions where heavy metal accumulation in the environment is a result from lack of treatment facilities that support proper management of waste and other related activities that provide fruitful opportunities for co-resistance to occur over prolonged periods of time^13^.

Given the complex matrix of the wastewater environment, robust *in vitro* investigation can help us to better understand which components (or which combination of components, or what concentrations of components) promote AMR. However, one critical gap is that current laboratory studies, which serve as evidence for resistance development, are typically performed in rich media. Opposed to rich media, it has been shown that bacteria grown in water systems have altered characteristics, such as cell envelope composition and morphology^14-16^. In addition, nutrient limitations present in wastewater can serve as environmental stressors to develop AMR^17^. Thus, experiments and models with rich media may not be a good representation of how AMR develops in wastewater. Currently, there are a few standard synthetic wastewater recipes which are used for studies on wastewater processing^18-21^. However, these have not been applied widely to bacterial or AMR studies. Moreover, there is limited evidence on how bacteria behave in real wastewater samples^22^. Thus, we seek to characterize bacterial growth, antibiotic susceptibility, and antibiotic resistance development in standard rich media, synthetic wastewater and real wastewater.

## Methods

### Strains and media conditions

*E. coli* MG1655 (ATCC 700926) was used for all experiments. LB broth was used for the control media. Synthetic wastewater was composed according to OECD guidelines with the components listed in Table 1 mixed in 1L of DI water^18,19^.

**Table 1.** Components and quantities of synthetic wastewater per 1L DI water^18,19^.

| Component | Quantity |
| --- | --- |
| Peptone | 160 mg |
| Meat extract | 110 mg |
| Urea | 30 mg |
| Anhydrous dipotassium hydrogen phosphate (K<sub>2</sub>HPO<sub>4</sub>) | 28 mg |
| Sodium chloride (NaCl) | 7 mg |
| Calcium chloride dihydrate (CaCl<sub>2</sub>·2H<sub>2</sub>O) | 4 mg |
| Magnesium sulfate heptahydrate (MgSO<sub>4</sub>·7H<sub>2</sub>O) | 2 mg |

### Wastewater sampling

Wastewater “influent” samples from the Massachusetts Water Resource Authority (MWRA) arrive at the plant through four underground tunnels. DITP pumps and lifts the influent from 80 to 150 feet to the head of the plant. There are three main pump stations at the plant. The 24 four composite influent collected was pumped into two two-liter bottles and sealed for pick up.

Following the collection of the influent, the rest of the samples undergo primary treatment. Grit is removed and disposed of at landfills and clarifiers are used to remove pollutants. At this stage 60% of suspended solids and 50% of pathogens and toxic chemicals are removed. The wastewater moves to the secondary treatment where mixers, reactors and clarifications remove remaining solids through biological and gravitational processes. Deer Island manufactures oxygen to feed microorganisms which consume dissolved organic matter and lead to 85% of pollution being removed from the wastewater^23^. Following this procedure, wastewater “effluent” is pumped into two-two liter bottles.

Samples were then collected from the WWTP on March 7, 2024, and brought to the autoclave on a liquid cycle at 121°C for 45 minutes less than an hour after being sampled.

### Bacterial growth in synthetic wastewater and autoclaved wastewater samples

To monitor growth, O.D. 600 was measured every 5 minutes for 48 hours using a Biotek plate reader with shaking in between each measurement. Wells were seeded with exponential phase Wild type E. coli MG1655 such that the starting O.D. 600 of each well was ∼0.08–0.09. To avoid condensation at 37 °C we made the plate cover hydrophilic as previously described^24^.

To measure viable cell growth, wild-type *E. coli* MG1655 was cultured at 37°C in 4mL of the media of interest in culture tubes and sampled at 0, 1 and 24 hours. These samples were plated in triplicate on LB agar to determine the CFU/mL at the time point of interest.

### MIC in synthetic wastewater and autoclaved wastewater samples

Wild-type *E. coli* MG1655 cultured at 37°C in the media of interest was grown in 96-well plates with 2-fold increments of ciprofloxacin, doxycycline, ampicillin and erythromycin. Each media condition was run in biological triplicate (n=3). MIC was determined to be at the highest concentration of antibiotic, where no growth was observed.

### Rate of resistance development of E.coli in synthetic wastewater and autoclaved wastewater samples

Wild-type *E. coli* MG1655 cultured at 37°C in the media of interest was grown in 96-well plates with 2-fold increments of ciprofloxacin or doxycycline. Each media condition was run in biological triplicate (n=3). We selected the bacteria in the well closest to 50% of the inhibitory concentration (IC50) to seed new bacterial cultures on the same dose series of ciprofloxacin at ∼ 24 hours, for 10 days. Bacteria serially passaged in LB broth media for the duration of the experiments served as the control groups.

### Characterization of Wastewater Samples

Iron content of each wastewater sample was tested using Bartovation Iron test strips for free soluble Fe^2+^/Fe^3+^. pH of each wastewater sample was tested using Cytiva pH test strips. Organic matter and ammonium content were measured using COD digestion vials (kit 2125825, Hach Company) and N-Ammonia Reagent Set (kit 2606945, Hach Company) test kits, and concentrations were measured with a Hach DR900 Colorimeter (Hach Company, Loveland, CO, US).

## Results

### Characterization of different media and wastewater

We characterized relevant properties of LB, synthetic wastewater and autoclaved wastewater samples, including pH and important macro and micronutrients. The pH, the concentration of organic matter–measured as COD– and ammonium concentration for both real wastewater samples, synthetic wastewater and LB media can be observed in Table 2. The synthetic wastewater sample exhibited the lowest ammonium concentration, which could potentially limit growth. On the contrary, LB media contained the highest concentration of ammonium and organic matter, providing optimal nutrient conditions for robust growth. The influent and effluent autoclaved wastewater samples showed average concentrations of both ammonium and COD, comparable to other mainstream wastewater sources, which contain between 20 - 60mgN/L of ammonia and typical concentrations of COD below 1000 mg/L^25, 26^. Additionally, None of the real wastewater samples had an iron content within the detectable range of 0-100 ppm, which was consistent with both the synthetic wastewater and LB media.

**Table 2.** Sample characterization.

|  | Autoclaved<br>Wastewater -<br>Influent | Autoclaved<br>Wastewater -<br>Effluent | Synthetic<br>Wastewater (1x) | LB |
| --- | --- | --- | --- | --- |
| pH | 8 | 8 | 6 | 7 |
| COD (mg/L) | 155 | 43 | 289 | 16,600 |
| NH<sub>3</sub> (mg-N/L) | 30 | 19 | 5 | 84 |

### *E. coli* display reduced growth in synthetic and real wastewater

In order to compare the growth of *E. coli* in the synthetic wastewater recipes and in autoclaved wastewater samples to growth in LB broth, a standard nutrient-rich media, we performed plate reader growth curve measurements (Figure 1) over 48 hours. We observed reduced growth in the synthetic wastewater recipes, with the 25 times increased concentration of the literature synthetic wastewater recipe (25X) showing more growth than the 5 times increased concentration of the literature synthetic wastewater recipe (5X). The autoclaved wastewater samples also showed reduced growth compared to both the LB broth and the synthetic wastewater recipes. We also verified these growth curves with colony counts on LB agar plates after 1 hour and 24 hours (Figure 2). After 1 hour, all media conditions displayed similar *E. coli* concentrations. However, after 24 hours, *E. coli* colony counts reflected concentrations similar to that predicted by the plate reader growth curves. The relative growth rates to the WT (numerical value of 1, dashed line) were also reported. While all samples were statistically different from the WT, the effluent was statistically different from the influent, 25X Synthetic wastewater, and 5X synthetic wastewater. This indicates the synthetic wastewaters tested propagated growth more like the influent than the effluent and can be used as substitutes for LB.

**Figure 1.**
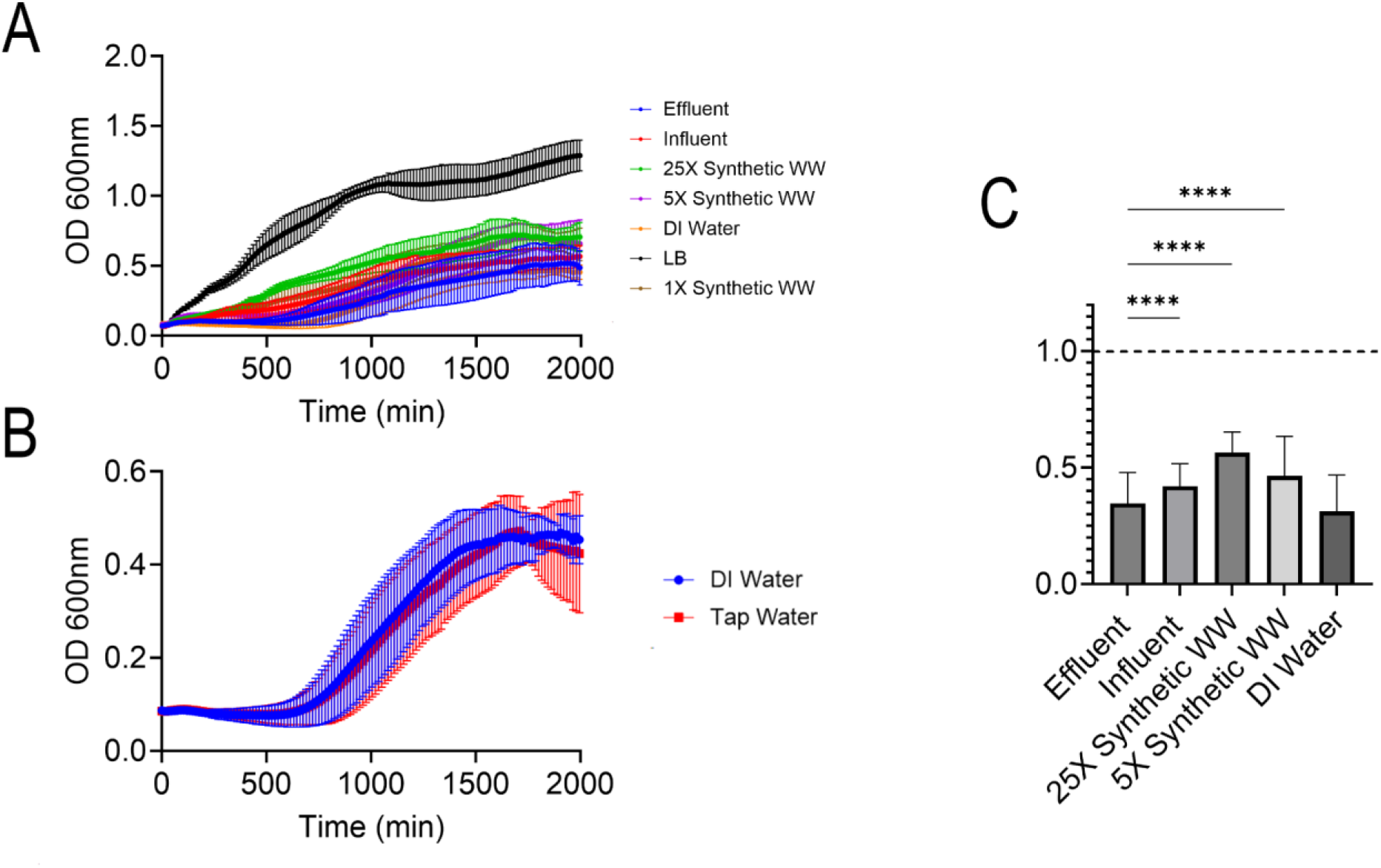
OD600 measurements of *E.coli* growth over 48 hours at 37C (A) comparison of influent and effluent autoclaved WW, 1X, 5X, and 25X Synthetic WW (SWW), DI Water and LB, (B) Autoclaved tap and (C) DI water and relative growth rates, **** represents population STDEV statistical significance.

**Figure 2.**
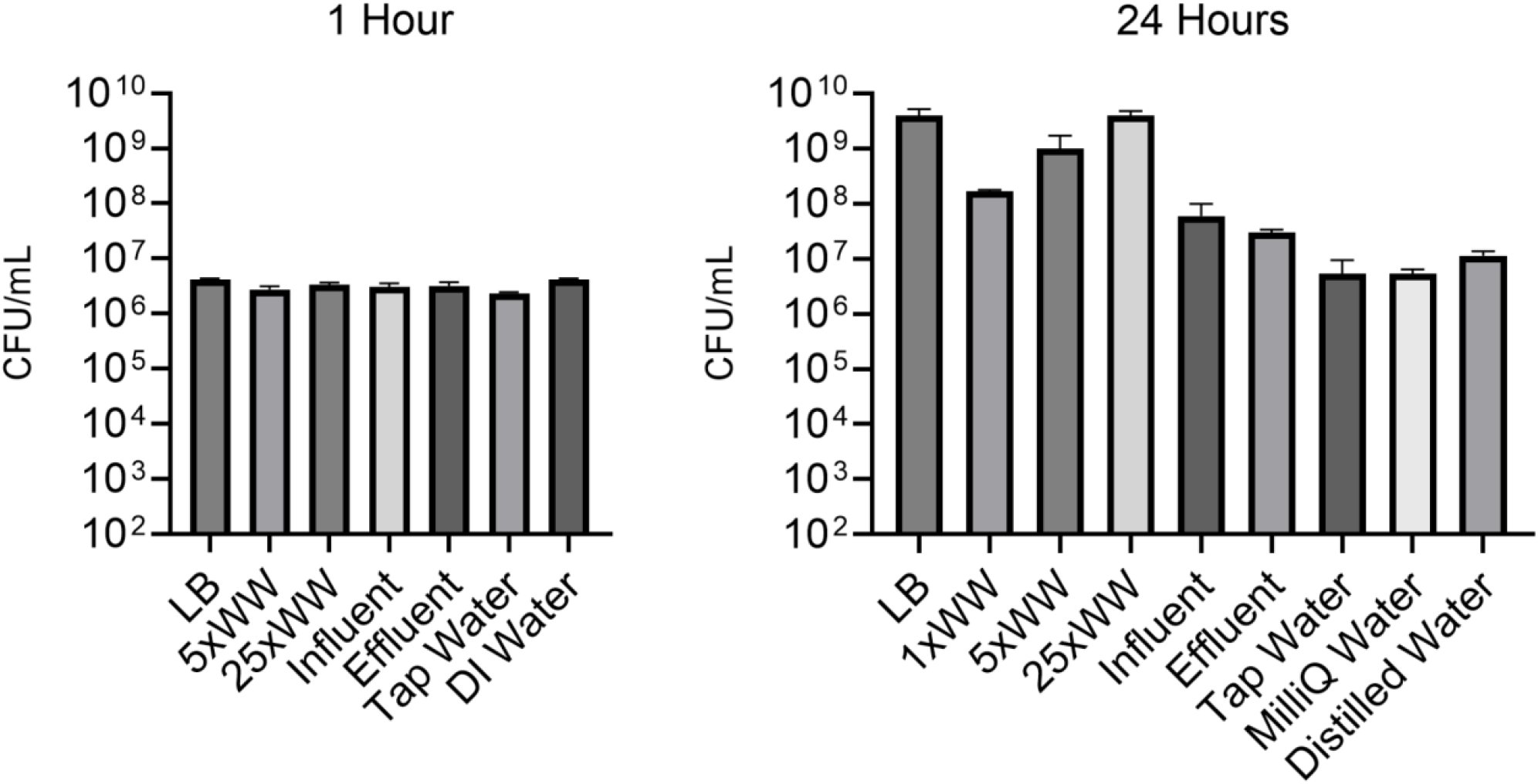
*E.coli* Colony counts on Agar Plates after 1 hour and after 24 hours

### E.coli grown in wastewater have altered sensitivities to antibiotics, suggesting synergistic and antagonistic relationships with wastewater components

In order to further compare the synthetic wastewater and the wastewater samples to each other and to LB media, we studied antibiotic activity against *E. coli* in each of these media. We first compared the MIC of doxycycline, an antibiotic that has been previously detected in wastewater, for *E. coli* in each of the media conditions (Figure 3). While similar MICs to doxycycline were observed in the synthetic wastewater and LB, the influent and effluent exhibited a significantly increased MIC. As one difference we noted among the different media was pH, we tested compositions with increased pH to match the basic pH of the real wastewater, and observed that the synthetic wastewater with a pH of 10 achieved a similarly increased MIC to the influent and effluent samples. However, the changed MIC in these wastewater samples and synthetic wastewater was not stable and reverted back to the wild-type MIC value when passaged for 24 hours in LB. This suggests that different characteristics and components of wastewater have synergistic and antagonistic interactions with antibiotics.

**Figure 3.**
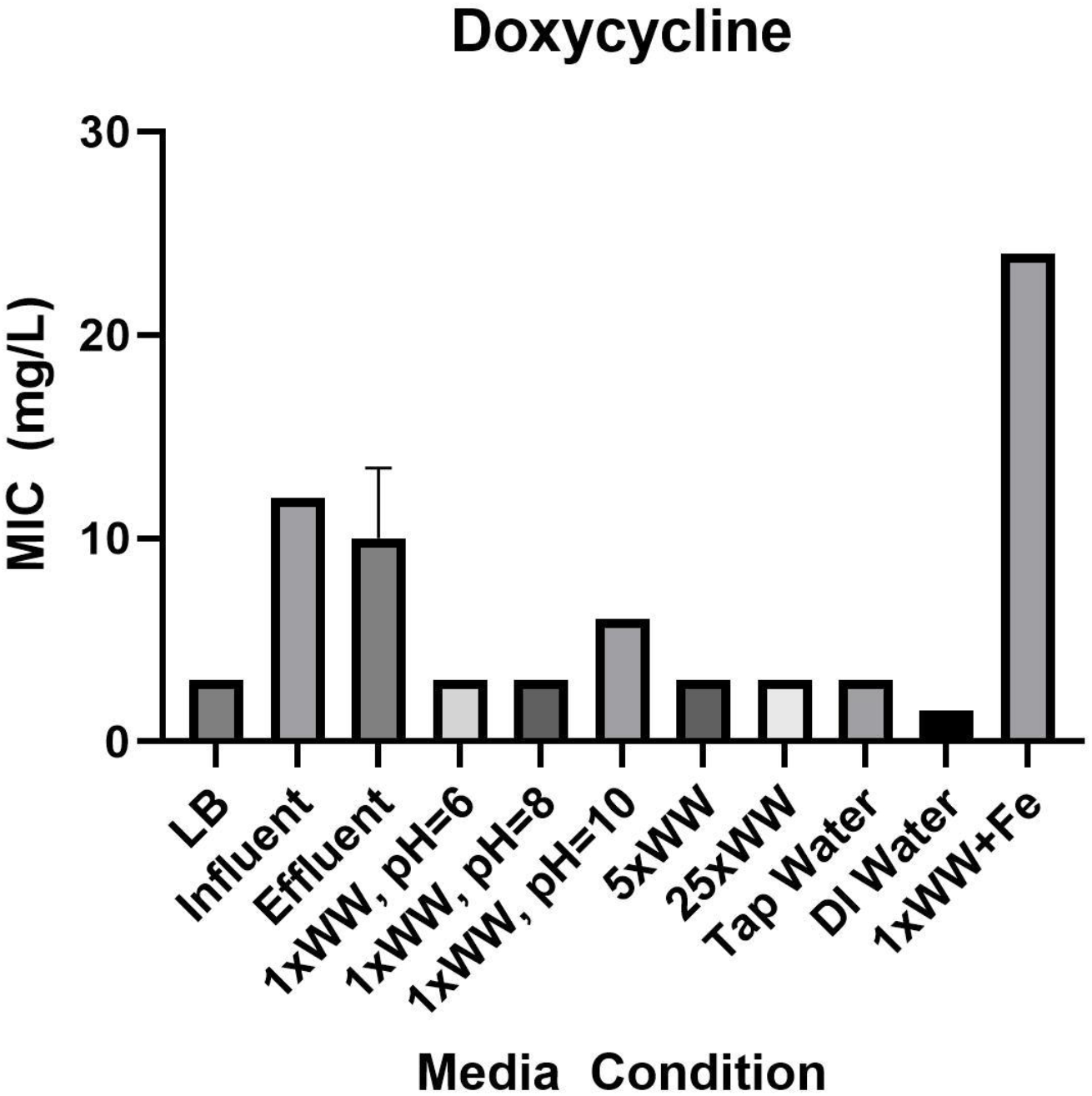
Comparison of Doxycycline MICs for various media conditions, error bars represent standard deviation

We next compared the MIC of ciprofloxacin for each of the media conditions (Figure 4). In this case, the less concentrated synthetic wastewater recipes exhibited decreased MICs compared to their respective decreased bacterial concentrations. This decreased MIC was also observed for the influent and effluent samples. We performed the same MIC assay with the synthetic wastewater at various pHs and with the real wastewater for Streptomycin, showing reduced MICs at high pHs and in the influent and effluent samples of real wastewater (Figure 5). As before, changes in MIC were not retained after passaged for 24 hours in LB.

**Figure 4.**
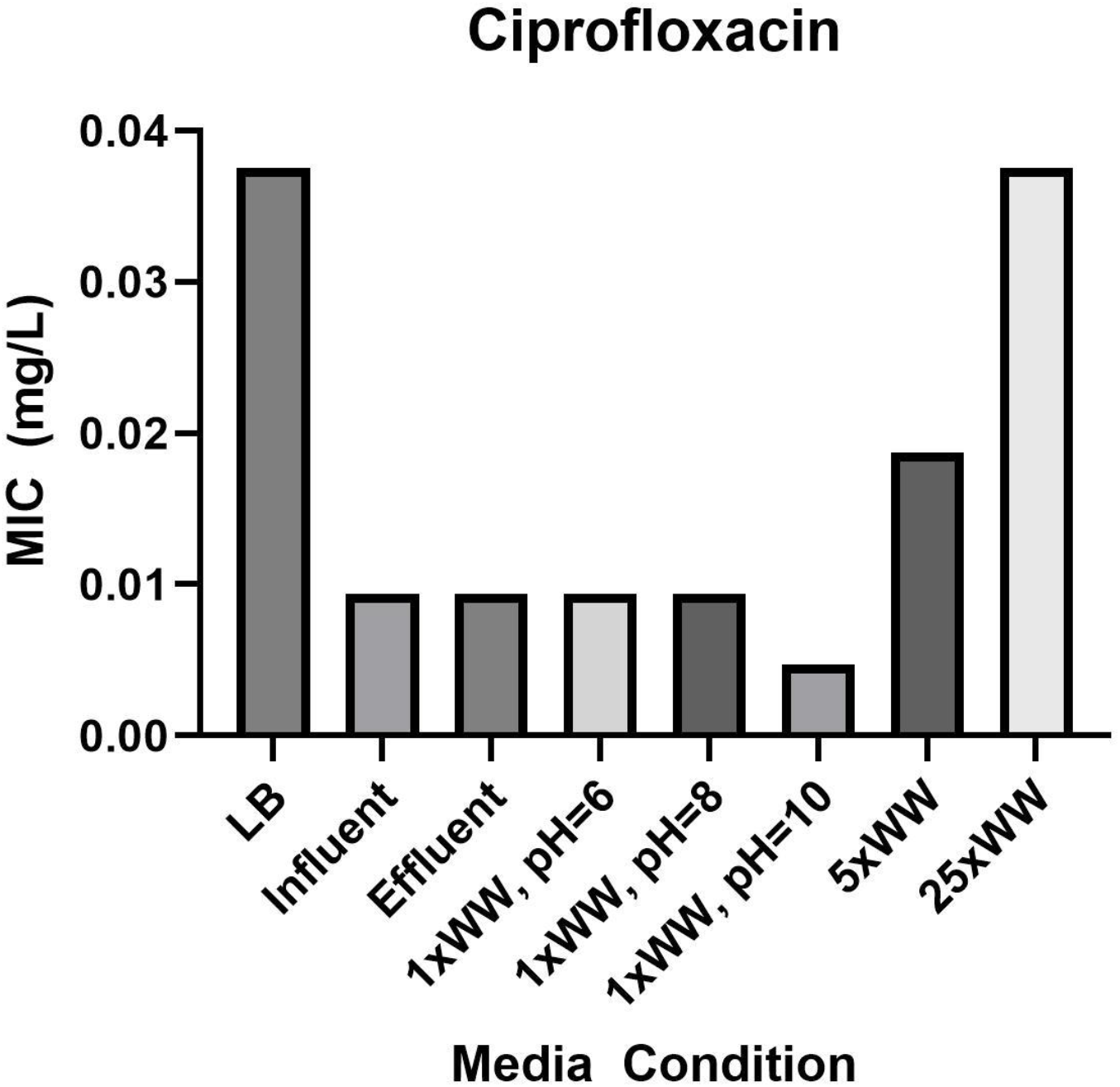
Comparison of Ciprofloxacin MICs for various media conditions, error bars represent standard deviation

**Figure 5.**
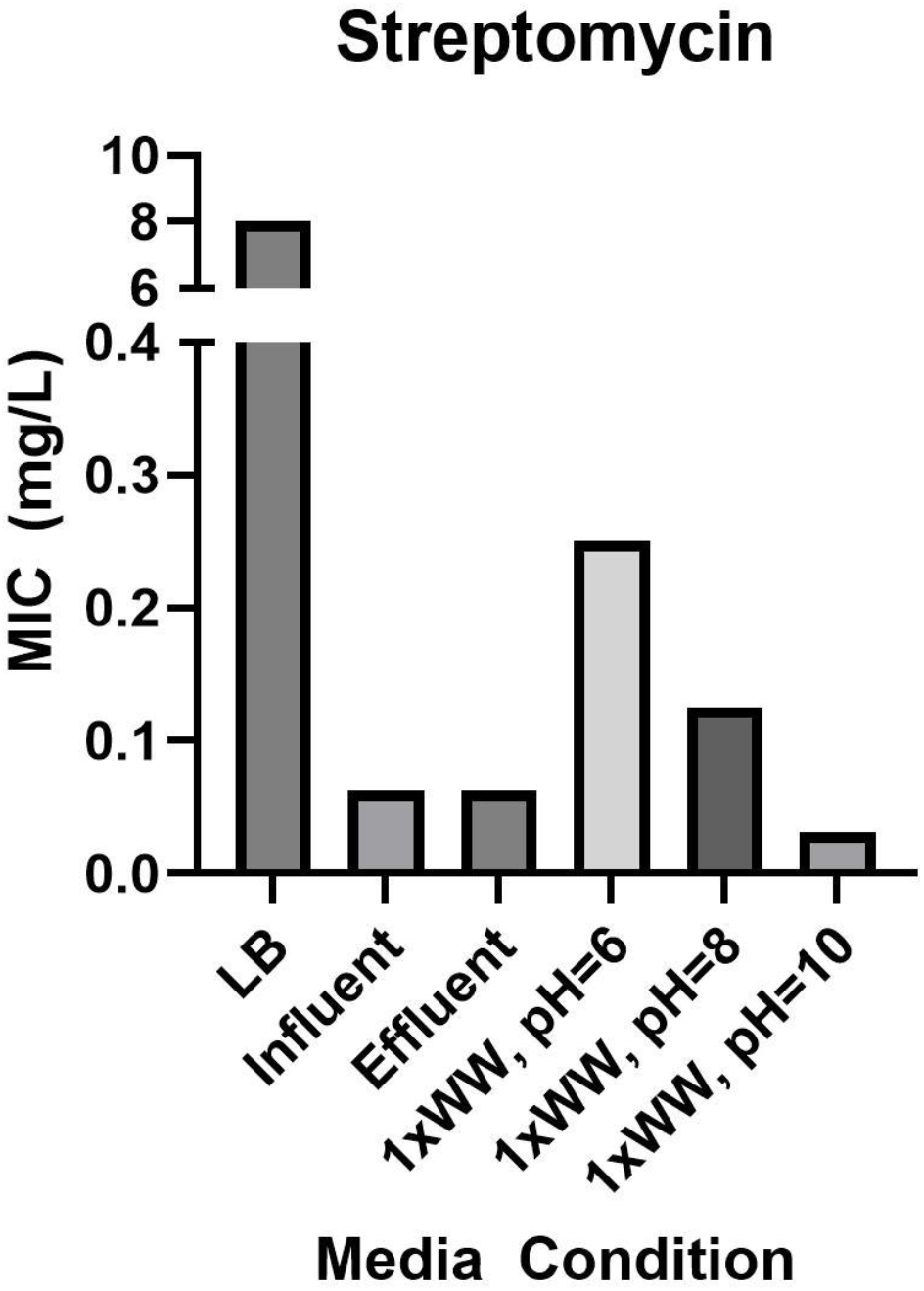
Comparison of Streptomycin MICs for various media conditions, error bars represent standard deviation

### Resistance develops in all medias, including MWRA wastewater samples, on similar timescales, but absolute values change based on different starting MICs, Iron speeds up resistance and leads to increased resistance

To further investigate the differences in antibiotic activity in the synthetic wastewater recipes and the wastewater samples as compared to each other and to LB media, we performed serial passaging experiments to probe how the different media affect the development of antibiotic resistance. Performing serial passaging in doxycycline (Figure 6) showed that while *E. coli* passaged in influent and effluent from the MWRA did not exhibit a high fold change in MIC over 10 days, the absolute MIC of all conditions not supplemented with iron developed to similar levels. Additionally, the supplementation of iron resulted in increased resistance in all of the media conditions. As iron is a metal pollutant of interest with interactions with common antibiotic residues, the validity of these interactions in wastewater and wastewater-like media is important to understanding AMR development in these environments.

**Figure 6.**
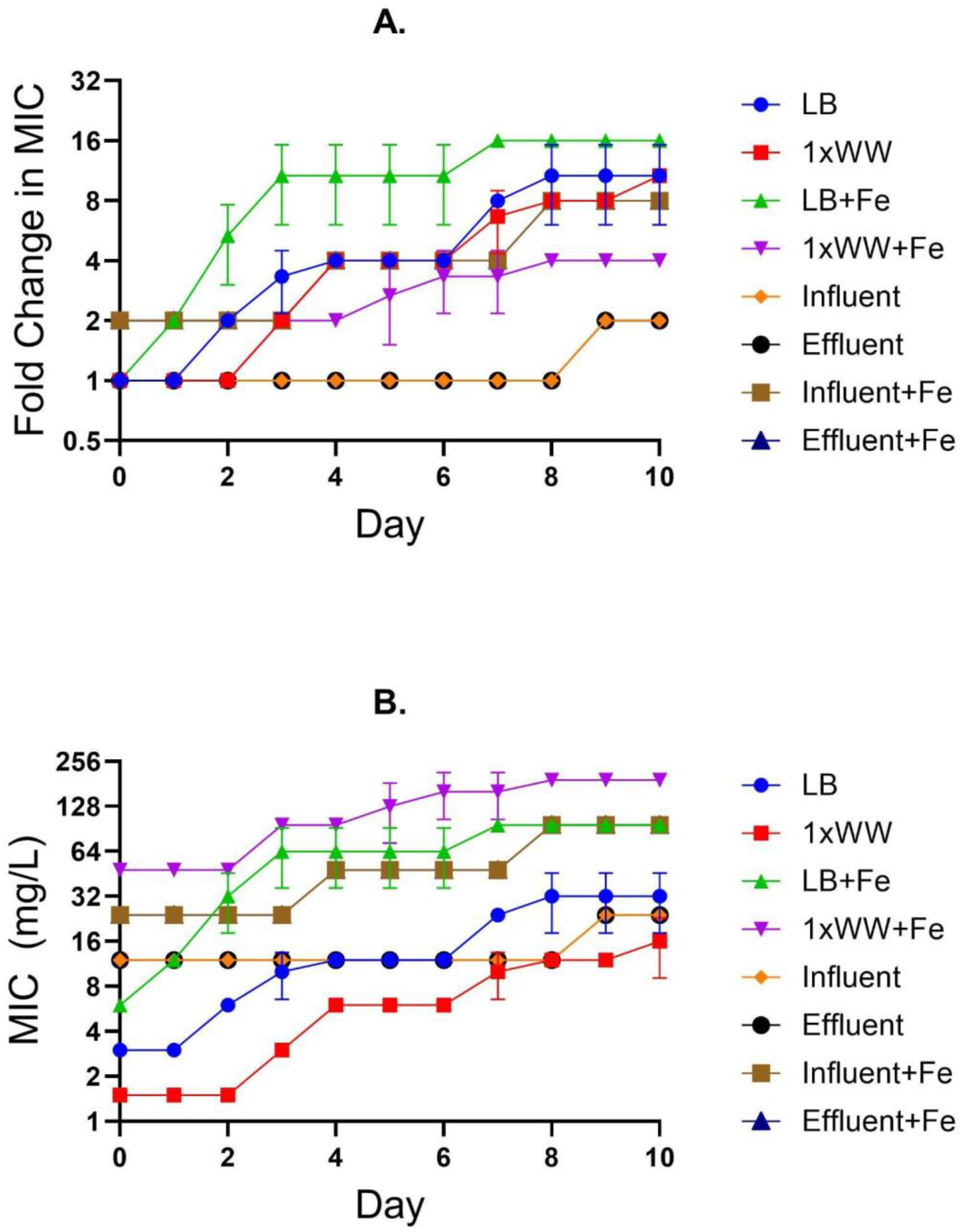
Change in Doxycycline MIC when passaged over time under different media conditions. **a.)** Fold change in Doxycycline MIC from baseline at Day 0. **b.)** Absolute MIC to Doxycycline over time

Performing serial passaging in ciprofloxacin (Figure 7) also exhibited similar absolute MIC values at the end of 10 days of serial passaging for the conditions not supplemented with iron. Furthermore, as observed in doxycycline, the media conditions supplemented with iron exhibited increased resistance development over time.

**Figure 7.**
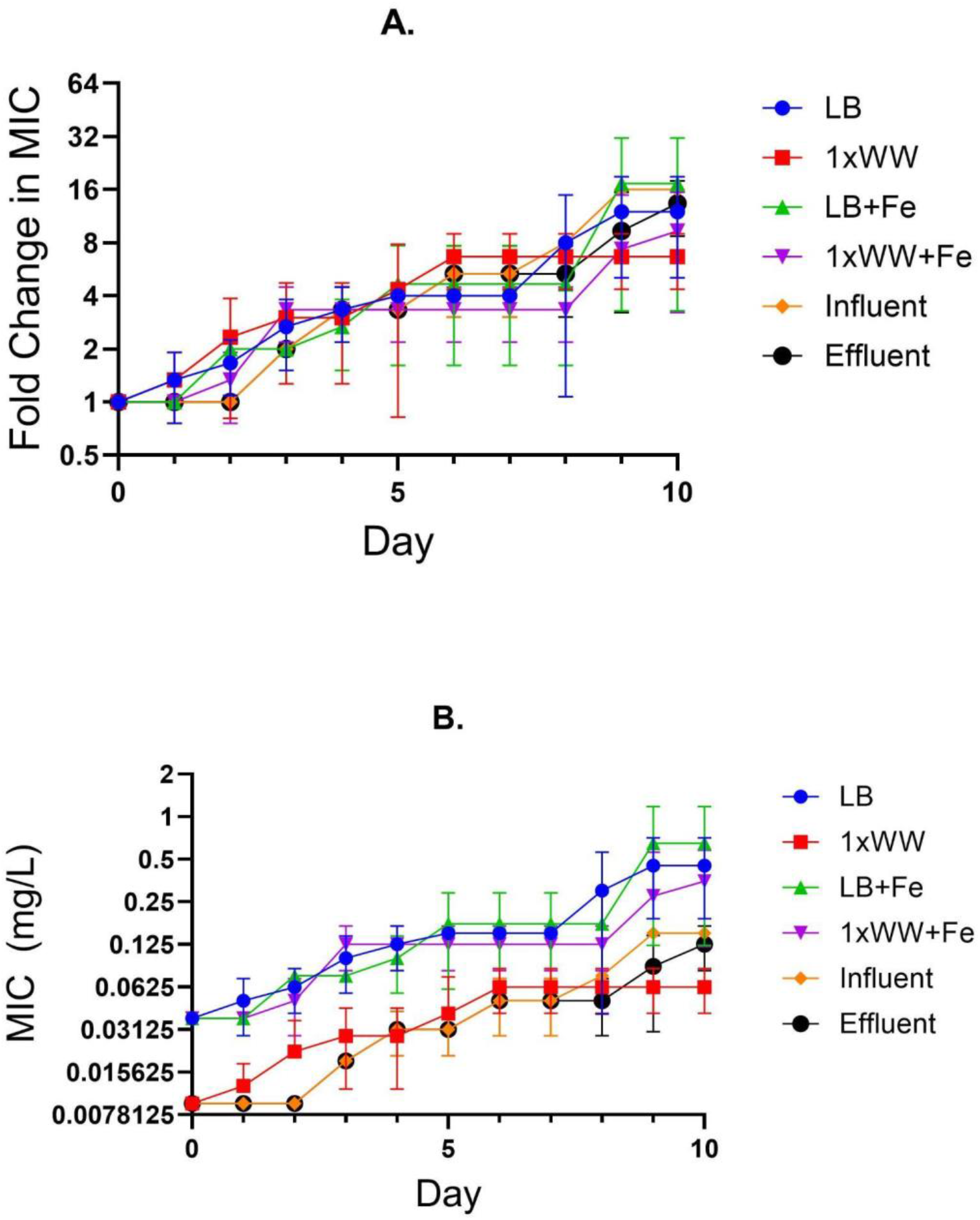
Change in Ciprofloxacin MIC when passaged over time under different media conditions. **a.)** Fold change in Ciprofloxacin MIC from baseline at Day 0. **b.)** Absolute MIC to Ciprofloxacin over time

We also wanted to probe the behavior of *E.coli* passaged in these media conditions with an antibiotic without a known interaction with iron, so we repeated the serial passaging experiment with streptomycin (Figure 8). In streptomycin, the media conditions all exhibit similar fold changes in MIC over the course of the experiment, with LB exhibiting a significantly higher absolute MIC and the synthetic wastewater and wastewater samples exhibiting similarly low absolute MICs.

**Figure 8.**
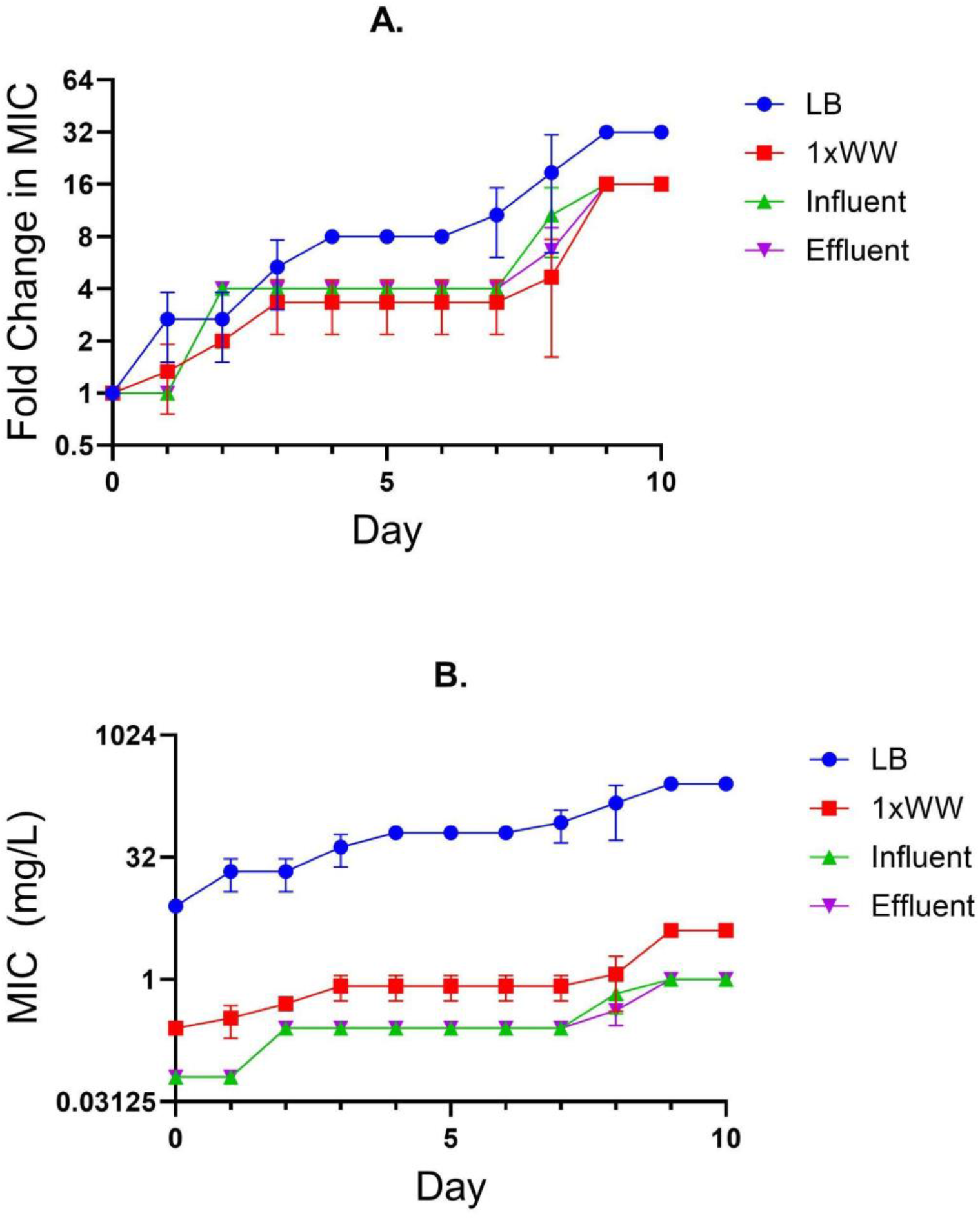
Change in Streptomycin MIC when passaged over time under different media conditions. **a.)** Fold change in Streptomycin MIC from baseline at Day 0. **b.)** Absolute MIC to Streptomycin over time

## Discussion

These studies in synthetic and real wastewater samples demonstrate that changes in media conditions can affect not only the growth of *E. coli*, but also the response of *E. coli* to antibiotic pressure. Both *E.coli* grown in synthetic wastewater and *E.coli* grown in autoclaved real wastewater showed reduced culture density after 24 hours compared to growth in LB, consistent with their reduced nutrient concentration. This effect was particularly pronounced in synthetic wastewater (1X), possibly due to the initial ammonium concentration–the most readily available nitrogen source–which was nearly at limiting concentrations from the start of the growth curve. In addition to the concentration of macronutrients, the varying concentrations of micronutrients in the real and synthetic wastewater samples–which were not measured in this study except for iron–may have influenced the growth rates of *E. coli*.

While the *E.coli* grown in synthetic wastewater showed reduced MICs to all antibiotics tested as compared to LB, the *E.coli* grown in autoclaved wastewater samples had a significantly higher MIC to Doxycycline, a comparable MIC to Ciprofloxacin and a significantly lower MIC to Streptomycin as compared to *E. coli* grown in LB media. This demonstrates that the standard media conditions used for *in vitro* studies of *E. coli* may yield results that are not fully representative of environmental *E. coli* behavior. Furthermore, differences in MIC in the real wastewater were not retained after 24 hours of passaging in LB, indicating that rather than affecting the bacteria, the wastewater had properties or components that were affecting the antibiotic activity in some way. One possibility is that the differing behavior was caused by the significant differences in pH of the synthetic wastewater and autoclaved MWRA wastewater when compared to LB media, with synthetic wastewater being more acidic than LB media and autoclaved wastewater being more basic than LB media. Prior literature has shown that pH higher than 8 can decrease Doxycycline absorption^27^, which is consistent with the decreased Doxycycline activity observed in the autoclaved real wastewater samples. Furthermore, *E. Coli* grown in basic compositions of synthetic wastewater were shown here to have similar susceptibility to Doxycycline, Ciprofloxaxcin and Streptomycin pressure to the *E. Coli* grown in the MWRA wastewater samples. Thus, we would suggest matching the pH of synthetic wastewater to the real wastewater samples of interest in order to better capture the effects of antibiotics on bacterial growth in these environments.

Additionally, we show resistance development under subinhibitory antibiotic pressure in both the synthetic and autoclaved real wastewater. While the absolute MICs of the antibiotics used differed between *E.coli* grown in the different media conditions over time, the studies in both Ciprofloxacin and Streptomycin showed similar fold changes in MIC over time as compared to the initial MICs in each media condition. This indicates that while there are differences in *E. coli* behavior in the synthetic wastewater and autoclaved real wastewater as compared to LB media, LB may be a sufficient proxy for these media when studying antibiotic resistance development under subinhibitory antibiotic pressure. While the case for Doxycycline is more complex, with *E. coli* developing far lower fold changes in MIC when serial passages in the autoclaved wastewater samples, the absolute MICs at the end of the serial passaging experiment were similar across the media conditions not supplemented with iron. This suggests that there may exist a maximum MIC to Doxycycline in the absence of iron that is conserved across the media conditions. The iron-supplemented synthetic wastewater, while exhibiting a similarly high starting MIC to Doxycycline as the autoclaved real wastewater, developed significantly higher Doxycycline resistance by the end of the experiment, thus making it an unsuitable proxy for real wastewater. However, the *E.coli* grown in autoclaved wastewater supplemented with iron still showed similar behavior to those grown in synthetic wastewater and LB media supplemented with iron, thus supporting the continued use of LB media in antibiotic resistance studies for *E. coli*.

One limitation of this study was the necessity of autoclaving the wastewater samples for use as a growth medium, which had the potential to degrade or alter some of the wastewater components. The experimental design required the use of sterile media to be comparable to lab made media such as LB. While we chose to autoclave the wastewater samples to maintain the larger solid particulate matter, filter sterilization could be used in future studies to better understand the composition of real wastewater. We were also limited in the number of wastewater samples tested here, with only influent and effluent samples from the MWRA in Massachusetts. Further studies with samples from other geographic locations nationally and internationally could further validate the findings here and shed light on key differences between wastewater environments across different treatment conditions and climates. Additionally, real wastewater environments typically have more than one bacterial species, and future studies with multiple bacterial species could further elucidate the validity of LB media and synthetic wastewater as proxies for real wastewater in *in vitro* studies. Further characterization of real wastewater samples would also provide greater insight into developing synthetic wastewater recipes that more closely recapitulate environmental wastewater conditions.

## Conclusion

Wastewater environments play a critical role in the development and spread of AMR, as resistant bacterial populations are frequently observed in these settings. However, most in vitro studies on AMR are conducted under media conditions that differ significantly from the composition and nutrient density found in actual wastewater. In our study, we observed that the use of synthetic wastewater and real wastewater samples as media for *in vitro* assays with *E. coli* resulted in differences in growth and response to antibiotic pressure as compared to LB broth. While the decreased growth in synthetic and real wastewater was correlated with decreased nutrient density in these media, differences in antibiotic susceptibility varied depending on the antibiotic used. However, when observing antibiotic resistance development under selective pressure from subinhibitory antibiotics and in the presence of environmental levels of iron, LB and synthetic wastewater captured the behavior of real wastewater to a significant degree, thus encouraging their use as a proxy for real wastewater in *in vitro* antibiotic resistance studies. These findings are crucial for enhancing public health surveillance methodologies, such as wastewater surveillance, as they offer valuable insights into analyzing the spread of AMR in different communities. Effective detection and monitoring of AMR can help develop public health strategies and interventions to prevent and control AMR.

## Acknowledgments

Authors would like to thank the Massachusetts Water Resources Authority for their assistance in procuring samples from the Deer Island Treatment Plant.

## CRediT Statement

**Indorica Sutradhar:** Conceptualization, Methodology, Formal Analysis, Investigation, Writing-Original Draft; **Neila Gross:** Conceptualization, Methodology, Formal Analysis, Investigation, Writing-Original Draft; **Carly Ching:** Conceptualization, Methodology, Writing-Original Draft; **Yanina Nahum:** Methodology, Investigation, Writing-Original Draft; **Darash Desai:** Conceptualization, Methodology, Writing-Review & Editing; **Devin Bowes:** Conceptualization, Methodology, Writing-Review & Editing**; Muhammad H. Zaman:** Conceptualization, Writing-Review & Editing, Supervision, Funding acquisition

## Data Sharing

All data used for this study has been included in the manuscript or supplementary material.

